# An annotated draft genome for *Radix auricularia* (Gastropoda, Mollusca)

**DOI:** 10.1101/087254

**Authors:** Tilman Schell, Barbara Feldmeyer, Hanno Schmidt, Bastian Greshake, Oliver Tills, Manuela Truebano, Simon D. Rundle, Juraj Paule, Ingo Ebersberger, Markus Pfenninger

## Abstract

Molluscs are the second most species-rich phylum in the animal kingdom, yet only eleven genomes of this group have been published so far. Here, we present the draft genome sequence of the pulmonate freshwater snail *Radix auricularia*. Six whole genome shotgun libraries with different layouts were sequenced. The resulting assembly comprises 4,823 scaffolds with a cumulative length of 910 Mb and an overall read coverage of 72x. The assembly contains 94.6 % of a metazoan core gene collection, indicating an almost complete coverage of the coding fraction. The discrepancy of ~690 Mb compared to the estimated genome size of *R. auricularia* (1.6 Gb) results from a high repeat content of 70 % mainly comprising DNA transposons. The annotation of 17,338 protein coding genes was supported by the use of publicly-available transcriptome data. This draft will serve as starting point for further genomic and population genetic research in this scientifically important phylum.

## Introduction

Gastropods are one of the broadest distributed eukaryotic taxa, being present across ecosystems worldwide. They occupy a maximally diverse set of habitats ranging from the deep sea to the highest mountains and from deserts to the Arctic, and have evolved to a range of specific adaptions (Romero et al. 2015, 2016). However, as for molluscs in general, whose species richness is second only to the arthropods (Dunn & Ryan 2015), gastropods are highly underrepresented among publicly available genomes (Figure 1). To date, only eleven mollusc genome sequences - of which six are from gastropods - exist with varying qualities concerning contiguity and completeness (Table 1). Any additional genome sequence has therefore the potential to substantially increase the knowledge about molluscs in particular and animal genomics in general.

**Fig. 1.**
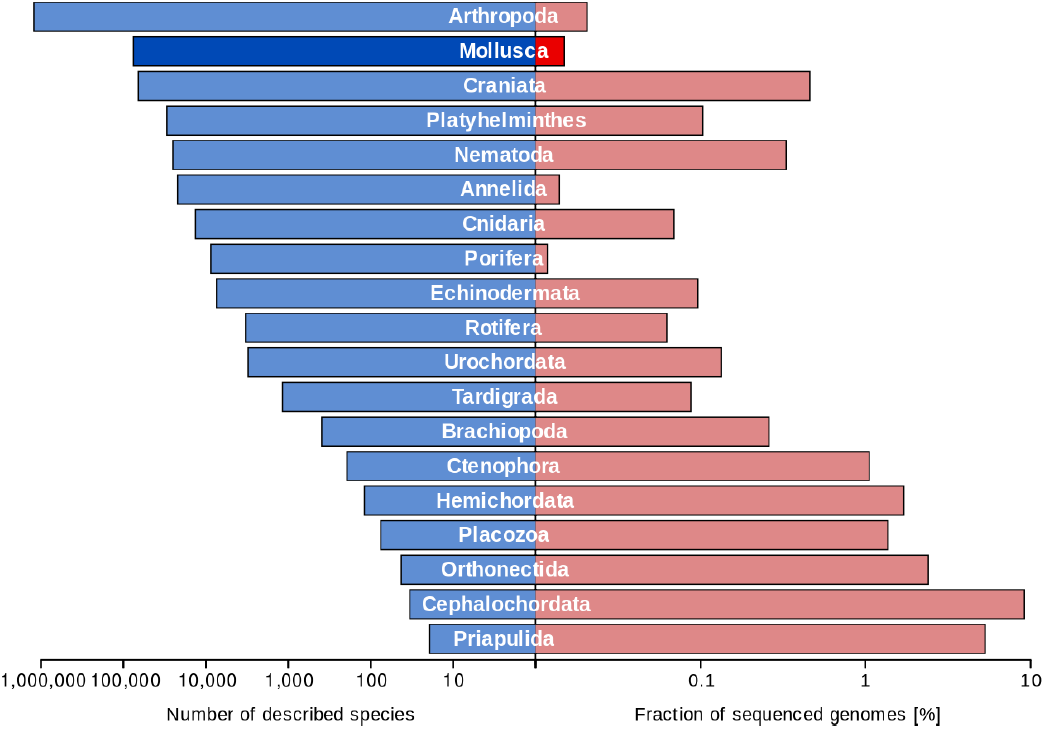
The number of described species (Dunn & Ryan 2015; GIGA Community of Scientists 2014) and the fraction of sequenced genomes (http://www.ncbi.nlm.nih.gov/genome/browse/ on September 1^st^, 2016). Animal phyla were obtained from (Dunn et al. 2014). Phyla with genomic record are displayed. Note the logarithmic scaling.

**Table 1.**
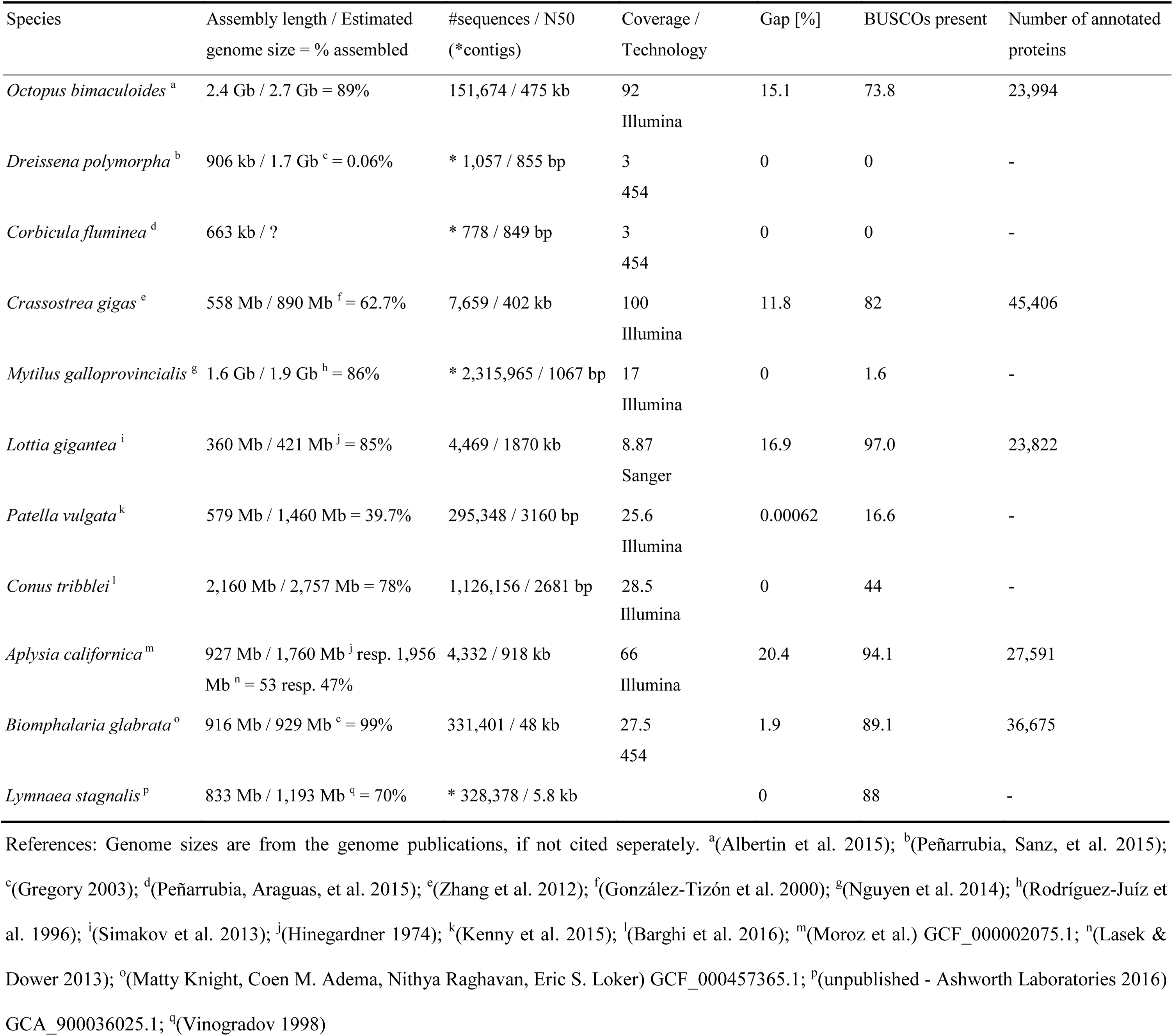
Available mollusc genomes. An overview from column 2 can be found in Supplementary Figure 4. Column 5: Fraction of N’s in the assembly. Column 6: BUSCOs: (Benchmarking Universal Single-Copy Orthologs) N_Metazoa_ = 843; Present = complete + fragmented.

The pulmonate freshwater snail genus *Radix* has a holarctic distribution (Glöer, Meier-Brook 1998; Cordellier et al. 2012) and plays an important role in investigating climate change effects in freshwater ecosystems (Sommer et al. 2012). The number of European species as well as the precise evolutionary relationships within the genus are controversial. This is mainly due to weak morphological differentiation and enormous environmental plasticity across species (Pfenninger et al. 2006). Members of the genus are simultaneously hermaphroditic (Yu et al. 2016; Jarne & Delay 1990) and both outcrossing and self-fertilisation occur (Jarne & Delay 1990; Jarne & Charlesworth 1993; Wiehn et al. 2002). The genus *Radix* is studied in many different fields, including parasitology (e.g. Huňová et al. 2012), evolutionary development (Tills et al. 2011), developmental plasticity (Rundle et al. 2011), ecotoxicology (e.g. Hallgren et al. 2012), climate change (Pfenninger et al. 2011), local adaptation (Quintela et al. 2014; Johansson et al. 2016), hybridisation (Patel et al. 2015) and biodiversity (Albrecht et al. 2012). Despite this broad range of interests, genomic resources are scarce and limited to transcriptomes (Feldmeyer et al. 2011, 2015; Tills et al. 2015) and mitochondrial genomes (Feldmeyer et al. 2010).

Here, we present the annotated draft genome sequence for *Radix auricularia* L. (Figure 2). This serves as an important foundation for future genomic and applied research in this scientifically important genus.

**Fig. 2.**
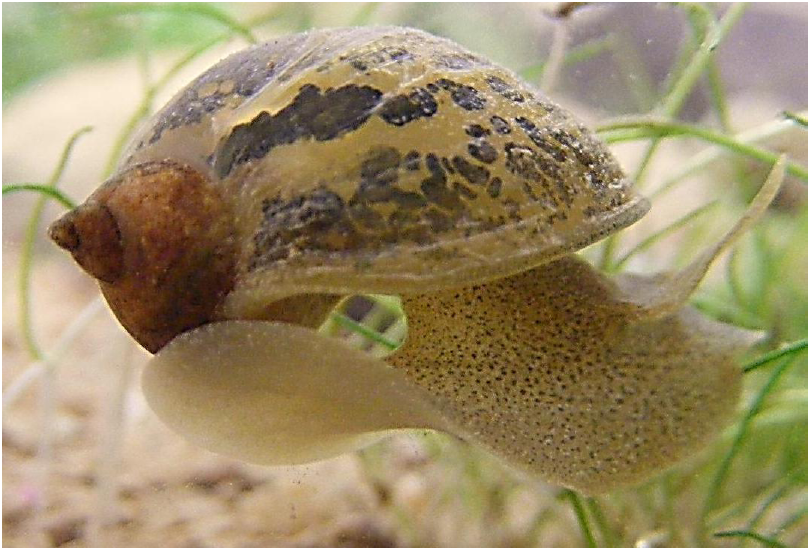
Photograph of *Radix auricularia*. Picture by Markus Pfenninger.

## Results and Discussion

### Genome assembly

A total of 1,000,372,010 raw reads (Supplementary Note 1; Supplementary Table 1) were generated and assembled into 4,823 scaffolds (Table 2; Supplementary Table 2). The mitochondrial genome (13,744 bp) was fully reconstructed, evidenced by comparison to the previously published sequence (Feldmeyer et al. 2015). Re-mapping the preprocessed reads revealed that 97.6 % could be unambiguously placed, resulting in a per position coverage distribution with its peak at 72x (Supplementary Note 2; Figure 3A). Additionally estimated insert sizes from mate pair libraries match their expected size (Figure 3B).

**Fig. 3.**
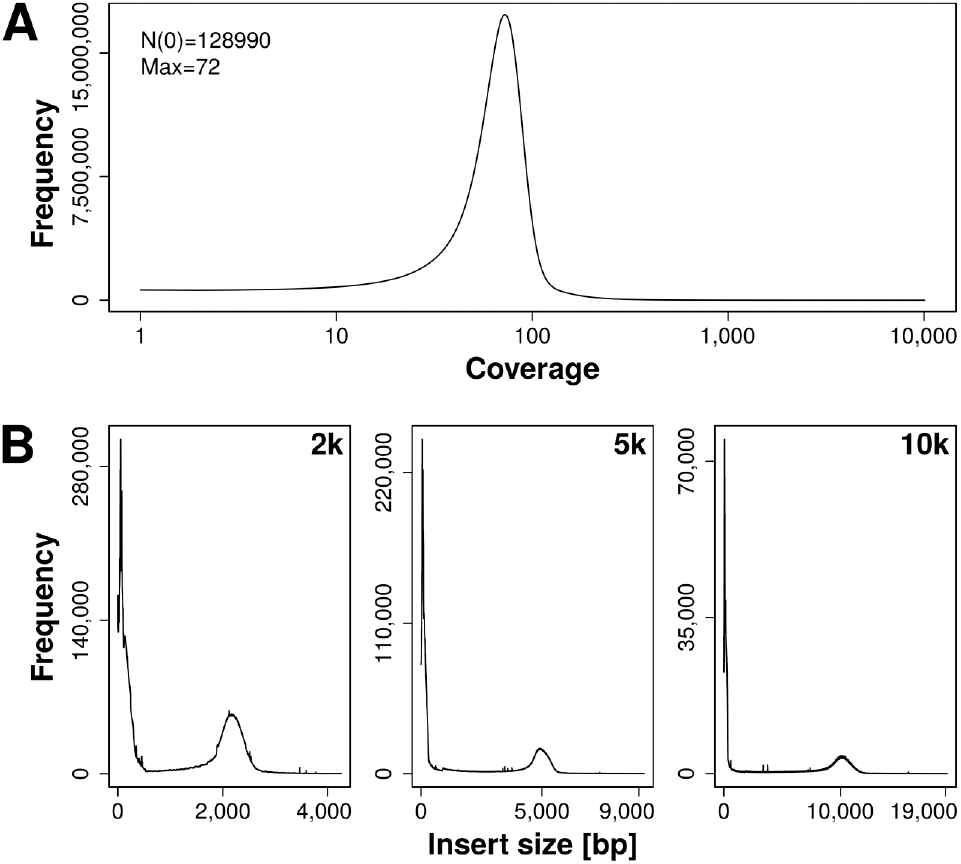
Re-mapping statistics. For details see Supplementary Note 2. A: Coverage distribution per position. The peak is located at a coverage of 72x. The x-axis is given in log-scale. B: Insert size distributions for the three mate pair libraries with insert sizes of 2, 5 and 10 kb. The high fraction mate pairs with insert sizes close to 0 (Supplementary Table 3) is due to the repetitive nature of the *Radix* genome. In particular repeat stretches that are not properly resolved in the genome assembly interfere with a proper placement of reads.

**Table 2.**
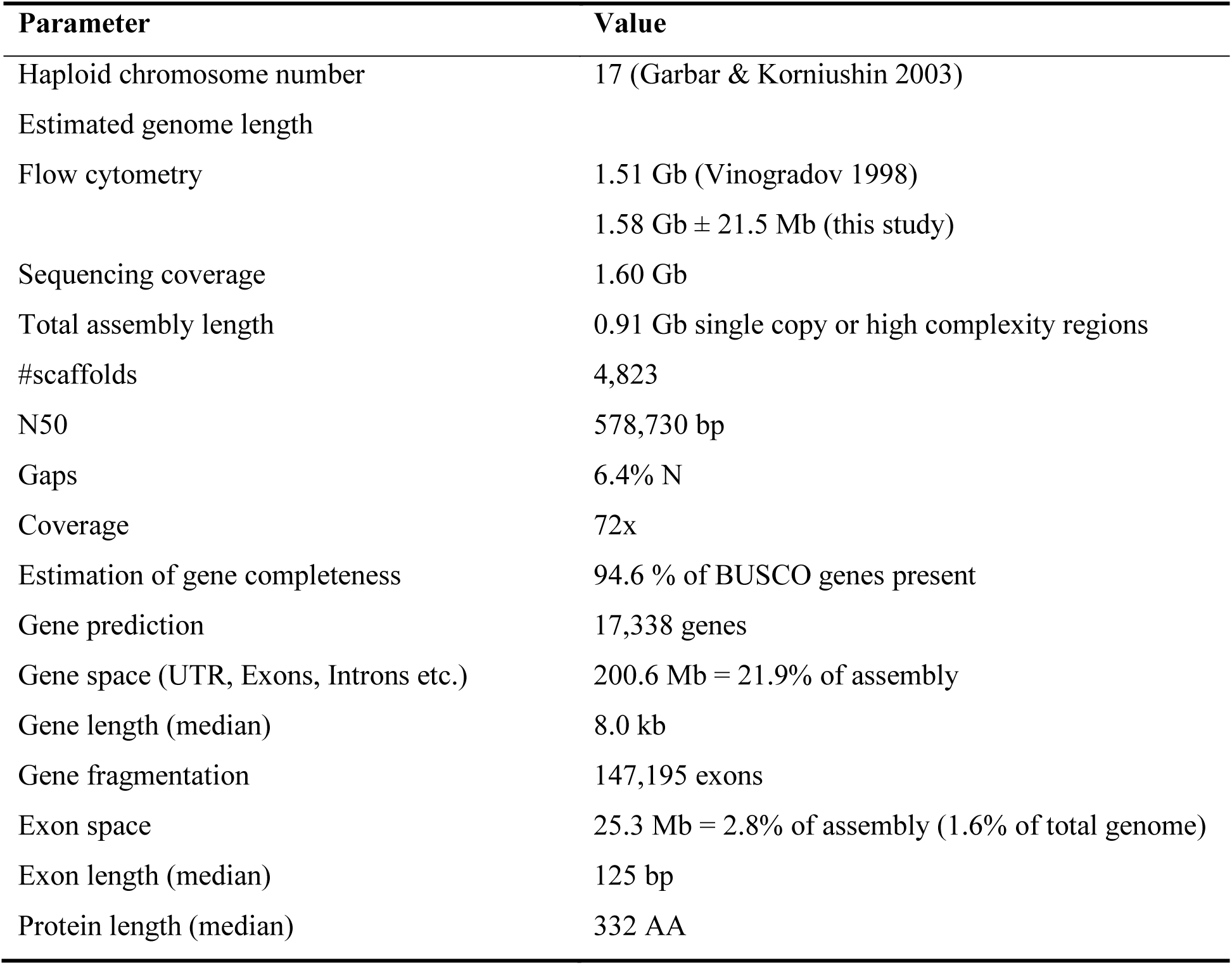
Characteristics of the *R. auricularia* genome and draft assembly. BUSCOs: (Benchmarking Universal Single-Copy Orthologs) N_Metazoa_ = 843; Present = complete + fragmented

The cumulative length of all scaffolds sums up to 910 Mb, which is about 665 Mb below the genome length estimates resulting from flow cytometric analyses (1,575 Mb; Supplementary Note 3) and from a read-mapping analysis (1,603 Mb; Supplementary Note 4). Both genome size estimates are consistent. This indicates an approximately uniform coverage of the nuclear genome in shotgun libraries without substantial bias introduced during library generation. This difference in length is most likely caused by a high repeat content in the *Radix* genome. Within scaffolds, 40.4 % of the sequence content was annotated as repeats mostly at the ends of contigs (Figure 4). This, in combination with a pronounced increase of read coverage at contig ends (Figure 4) is typical for collapsed repeat stretches. The overall repeat content of the genome was estimated to be approximately 70 % (Supplementary Note 5). The majority of repeats were either classified as Transposable Elements or as ‘unknown’ (Supplementary Figure 3). The difference between genome size and assembly length of this *R. auricularia* draft assembly resembles that of other published mollusc genomes (Supplementary Figure 4). However, when considering contiguity reflected in the N50 value it ranks among the top mollusc genomes (Table 1; Table 2). To evaluate completeness of the assembly’s gene space we used BUSCO (Simão et al. 2015) in combination with the provided metazoan set and recovered 94.6 % of the subsumed genes. This suggests no conspicuous lack of gene information.

**Fig. 4.**
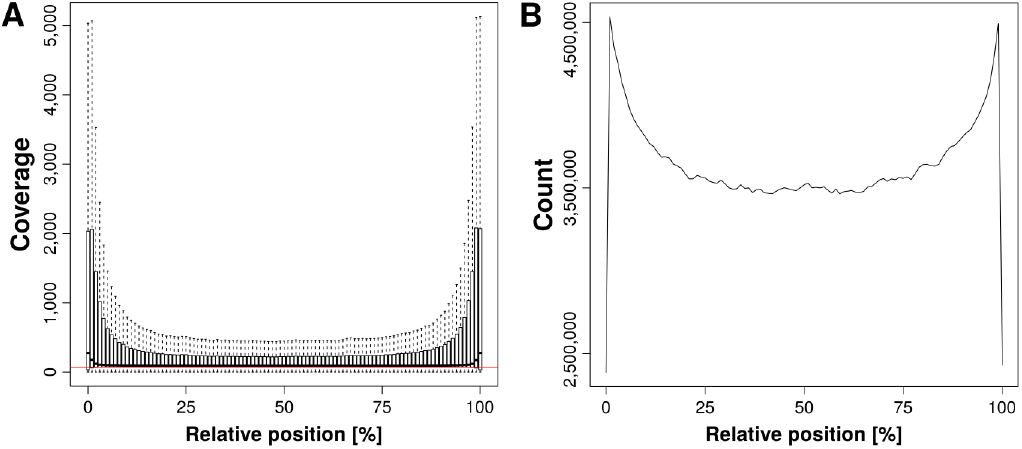
Collapsed repeats. A: Coverage of continuous unambiguous sequence parts of the scaffolds. Outliers from boxplots are not shown. The red line represents the most frequent coverage of 72x (Figure 2A). B: Positions annotated as repeats along continuous unambiguous sequence parts of the scaffolds.

### Genome Annotation

The annotation resulted in 17,338 protein coding genes (Table 2) of which 70.4 % show a significant sequence similarity to entries in the Swiss-Prot database (e-value < 10^-10^, accessed on May 11^th^ 2016).

The number of identified genes is at the lower end compared to other annotated mollusc genomes (Min: *Lottia gigantea* 23,822; Max: *Crassostrea gigas* 45,406; Table 1; Supplementary Table 4). Thus predicted *Radix* proteins were screened for completeness regarding evolutionary conserved genes using HaMStR (Ebersberger et al. 2009). The analysis resulted in a recovery of 93.7 % and is in line with the results from BUSCO. Extrapolating completeness estimates of both tools suggests that the annotation covers the majority of genes being present in the draft genome sequence. We then checked how the differences in protein numbers could be explained. The fraction of orthogroups (cluster of orthologous genes; see Material and Methods) containing only one sequence per species was highest in *Radix* (Supplementary Figure 5). Moreover there was a negative correlation (R^2^ = 0.77; p = 0.02) between the number of annotated proteins per species and fraction of orthogroups containing only one sequence per species (Supplementary Figure 6). One explanation for this observation could be lineage specific gene duplications in the other mollusc lineages. Additionally, artificial gene fissions in the course of genome annotation may be less common in *Radix*. This might be attributed to our use of comprehensive transcriptomic data of *Radix* for guiding gene prediction.

Next to the evolutionarily old genes represented in the BUSCO and HaMStR gene sets, *Radix* contains 1,481 genes for which we could find no orthologs in the other mollusc and additional non-mollusc spiralian gene sets (Supplementary Table 5; Supplementary Note 6). We tested for overrepresentation of functional categories in genes private to *Radix*, as well as in genes present in all molluscs but *Radix*. We identified 17 Gene Ontology (GO) terms to be significantly enriched amongst the 1,481 proteins private to *Radix* compared to all other mollusc and non-mollusc spiralian protein sets available. Enriched terms include “nucleoside transmembrane transport”, “carbohydrate metabolic process” and “chitin catabolic process” (Supplementary Table 6). Among the categories found in all annotated molluscs but *Radix* (Supplementary Table 7; Supplementary Note 6), the “G-protein coupled receptor signalling pathway” is the most prominent one. G-protein receptors are involved in reactions to “hormones, neurotransmitters and environmental stimulants” (Rosenbaum et al. 2009). The loss of these genes could have led to reduced sensitivity to such stimuli in *Radix*. Whether the reduced number of G-coupled receptor pathway components is biologically meaningful, or a result of technical and analytical limitations, cannot be determined from the present data. Membrane proteins, for example, are generally more diverse than water soluble proteins in the tree of life (Sojo et al. 2016), so we hypothesise that its proteins could be highly modified and were thus not identified as such in *Radix*.

### Conclusion

Here we present a draft genome of the snail *Radix auriculara*. The genome is comparable in size to other mollusc genomes and also rich in repeats. This new genomic resource will allow conducting future studies on genome evolution, population genomics and gene evolution within this genus and higher gastropod and mollusc taxa.

## Material and Methods

### Sample collection and sequencing

Snails were collected from a pond in the Taunus, Germany, identified with COI barcoding (Pfenninger et al. 2006) and kept under laboratory conditions for at least five generations of inbreeding by full-sib mating. Three specimens of *R. auricularia* (Figure 2) were used for DNA extraction. Pooled DNA was used for preparation of three paired end and three mate pair (2 kbp, 5 kbp, 10 kbp insert size) libraries, that were sequenced on an Illumina HiSeq 2000 and 2500 at Beijing Genomics Institute, Hong Kong (Supplementary Note 1; Supplementary Table 1). Reads were cleaned of adapter sequences using Trimmomatic 0.33 (Bolger et al. 2014; Supplementary Note 7) and screened for contaminations with FastqScreen 0.5.2 (http://www.bioinformatics.babraham.ac.uk/projects/fastq_screen/; Supplementary Note 8; Supplementary Figure 7). Raw reads have been deposited under NCBI BioProject PRJNA350764.

### Genome size estimation

Genome size was estimated by flow cytometry based on a modified protocol of (Otto 1990; Supplementary Note 9). Additionally, we estimated the genome size from our sequence data by dividing the total sum of nucleotides used for the assembly by the peak coverage from mapping back the assembly reads with the *bwa mem* algorithm from BWA 0.5.10 (Li 2013; Supplementary Note 4). Re-mappings were also used to estimate the repeat content of the genome (Supplementary Note 5).

### Assembly strategy

Reads were assembled using the Platanus 1.2.1 pipeline (Kajitani et al. 2014) with k-mer sizes ranging from 63 to 88 and a step size of 2. All other assembly parameters were kept at the default value. The output of the Platanus pipeline was filtered for sequences ≥ 500 bp. Afterwards, scaffolding was performed using SSPACE 3.0 (Boetzer et al. 2011) with ‘contig extension’ turned on. To further increase the contiguity of the draft genome we applied a third scaffolding step, making use of the cDNA sequence data. Transcriptome contig sequences of *R. auricularia* and three closely related species (Supplementary Note 10) were mapped sequentially according to phylogeny (Feldmeyer et al. 2015) using BLAT 35 (Kent 2002), with *-extendThroughN* enabled apart from default settings, onto the scaffolds; the gapped alignments were then used for joining of sequences with L_RNA_scaffolder (Xue et al. 2013). Finally, all sequences with at least 1,000 bp were used as input for GapFiller 1.10 (Boetzer et al. 2012) to close extant gaps in the draft genome. Details of the assembly can be found in Supplementary Note 11.

### Annotation strategy

Metazoan core orthologous genes were searched in the *R. auricularia* assembly and all other available mollusc genomes using BUSCO 1.2b (Simão et al. 2015).

The whole annotation process was performed using the MAKER2 2.31.8 pipeline and affiliated programs (Cantarel et al. 2008; Holt & Yandell 2011). Initially, we built a custom repeat library from the assembly using RepeatModeler 1.0.4 (Simit & Hubley 2015) and read data using dnaPipeTE 1.2 (Goubert et al. 2015) with 30 upstream trials on varying coverage depths and then 50 parallel runs on the best-fitting coverage of 0.025 (Supplementary Note 12). The draft genome and transcriptome of *R. auricularia* (Supplementary Note 10) in addition to the BUSCO 1.2b (Simão et al. 2015) annotations of core metazoan genes on the draft genome were used as input for the initial training at the Augustus webserver (Stanke et al. 2004; http://bioinf.uni-greifswald.de/webaugustus/). As additional input for MAKER2, we created two hidden Markov models on the gene structure of *R. auricularia*. One was generated by GeneMark 4.32 (Lomsadze et al. 2005) and another by SNAP 2006-07-28 (Korf 2004), using the output of CEGMA v2.5 (Parra et al. 2007; summarized results in Supplentary Table 9). We ran three consecutive iterations of MAKER2 with the draft genome sequence, the transcriptomes (Supplementary Note 10), models from Augustus, SNAP and GeneMark, the repeat library and the Swiss-Prot database (Accessed at May 23^th^ 2016). Between the iterations, the Augustus 3.2.2 (Stanke et al. 2004) and SNAP models were retrained according to the best-practice MAKER2 workflow (Supplementary Note 13). Finally, all protein sequences from MAKER2 output were assigned putative names by BLASTP searches (Camacho et al. 2009) against the Swiss-Prot database. In addition we used the targeted ortholog search tool, HaMStR v. 13.2.6 (Ebersberger et al. 2009; http://www.sourceforge.net/projects/hamstr/) to screen for 1,031 evolutionarily conserved genes that pre-date the split of animals and fungi. HaMStR was called with the options -strict, -checkcoorthologsref, and -hitlimit=5. The profile hidden Markov models that served as input for the search are included in the HaMStR distribution.

We created orthologous groups from protein sequences of all six annotated molluscs and 16 additional non-mollusc spiralian species with OrthoFinder 0.7.1 (Emms & Kelly 2015). All proteins were functionally annotated using InterProScan 5 (Quevillon et al. 2005; Zdobnov & Apweiler 2001). The enrichment analyses were performed in TopGO (Alexa & Rahnenfuhrer 2016), a bioconductor package for R (R Development Core Team 2008). We tested for significant enrichment of GO terms in proteins private to *Radix* and proteins found in all molluscs but *Radix*. We applied a Fischer’s exact test, FDR correction and filtered by q-values smaller than 0.05. Additional information can be found in Supplementary Note 6.

## Supplementary Material

Supplementary Note 1. Individuals and Sequencing

Supplementary Table 1. Sequenced raw data

Supplementary Table 2. Summary statistics of different assembly steps

Supplementary Note 2. Re-mapping

Supplementary Figure 1. Number of partially mapped reads along continuous parts of scaffolds

Supplementary Figure 2. Mapping quality frequency distribution

Supplementary Table 3. Re-mapping statistics from mate pair libraries

Supplementary Note 3. Results from flow cytometric measurements

Supplementary Note 4. Genome size estimation from coverage

Supplementary Note 5. Repeat content

Supplementary Figure 3. Classified repeat families

Supplementary Figure 4. Mollusc genome sizes

Supplementary Table 4. Proteins similar to Swiss-Prot entries

Supplementary Figure 5. Number of sequences from one species in orhtogroups

Supplementary Figure 6. Regression of protein sequences and orthogroups

Supplementary Table 5. Protein sets used for ortholog clustering and GO-term enrichment

Supplementary Note 6. Orthologous clustering and Gene Ontology enrichment

Supplementary Table 6. Enriched GO-terms for *Radix*

Supplementary Table 7. Enriched GO-terms not in *Radix*

Supplementary Note 7. Preprocessing and trimming

Supplementary Note 8. Contamination screening

Supplementary Figure 7. Results of contamination screening

Supplementary Note 9. Material and methods of flow cytometric analysis

Supplementary Note 10. Transcriptome assemblies

Supplementary Note 11. Genome assembly

Supplementary Figure 8. Transcriptome filtering

Supplementary Table 8. Genome scaffolding with transcriptomic data

Supplementary Note 12. Repeat library

Supplementary Figure 9. Read subsampling

Supplementary Table 9. Summarized results of the CEGMA analysis

Supplementary Note 13. Annotation

## Acknowledgments

The project was funded by SAW-network project 291. Thanks to Fabrizio Ghiselli and another anonymous reviewer for their helpful comments on the manuscript.

## Literature cited

Albrecht C, Hauffe T, Schreiber K, Wilke T. 2012. Mollusc biodiversity in a European ancient lake system: Lakes Prespa and Mikri Prespa in the Balkans. Hydrobiologia. 682:47–59. doi: 10.1007/s10750-011-0830-1.

Alexa A, Rahnenfuhrer J. 2016. topGO: Enrichment Analysis for Gene Ontology. R package version 2.26.0.

Boetzer M et al. 2012. Toward almost closed genomes with GapFiller. Genome Biol. 13:R56. doi: 10.1186/gb-2012-13-6-r56.

Boetzer M, Henkel C V., Jansen HJ, Butler D, Pirovano W. 2011. Scaffolding pre-assembled contigs using SSPACE. Bioinformatics. 27:578–579. doi: 10.1093/bioinformatics/btq683.

Bolger AM, Lohse M, Usadel B. 2014. Trimmomatic: A flexible trimmer for Illumina sequence data. Bioinformatics. doi: 10.1093/bioinformatics/btu170.

Camacho C et al. 2009. BLAST plus: architecture and applications. BMC Bioinformatics. 10:1. doi: Artn 421\nDoi 10.1186/1471-2105-10-421.

Cantarel BL et al. 2008. MAKER: An easy-to-use annotation pipeline designed for emerging model organism genomes. Genome Res. 18:188–196. doi: 10.1101/gr.6743907.

Dunn CW, Giribet G, Edgecombe GD, Hejnol A. 2014. Animal Phylogeny and Its Evolutionary Implications. Annu. Rev. Ecol. Evol. Syst. 45:371–395. doi: 10.1146/annurev-ecolsys-120213-091627.

Dunn CW, Ryan JF. 2015. The evolution of animal genomes. Curr. Opin. Genet. Dev. 35:25–32. doi: 10.1016/j.gde.2015.08.006.

Ebersberger I, Strauss S, von Haeseler A. 2009. HaMStR: profile hidden markov model based search for orthologs in ESTs. BMC Evol. Biol. 9. doi: 10.1186/1471-2148-9-157.

Emms DM, Kelly S. 2015. OrthoFinder: solving fundamental biases in whole genome comparisons dramatically improves orthogroup inference accuracy. Genome Biol. 16:157. doi: 10.1186/s13059-015-0721-2.

Feldmeyer B, Greshake B, Funke E, Ebersberger I, Pfenninger M. 2015. Positive selection in development and growth rate regulation genes involved in species divergence of the genus Radix. BMC Evol. Biol. 15:164. doi: 10.1186/s12862-015-0434-x.

Feldmeyer B, Hoffmeier K, Pfenninger M. 2010. The complete mitochondrial genome of Radix balthica (Pulmonata, Basommatophora), obtained by low coverage shot gun next generation sequencing. Mol. Phylogenet. Evol. 57:1329–1333. doi: 10.1016/j.ympev.2010.09.012.

Feldmeyer B, Wheat CW, Krezdorn N, Rotter B, Pfenninger M. 2011. Short read Illumina data for the de novo assembly of a non-model snail species transcriptome (Radix balthica, Basommatophora, Pulmonata), and a comparison of assembler performance. BMC Genomics. 12:317. doi: 10.1186/1471-2164-12-317.

GIGA Community of Scientists. 2014. The Global Invertebrate Genomics Alliance (GIGA): Developing Community Resources to Study Diverse Invertebrate Genomes. J. Hered. 105:1–18. doi: 10.1093/jhered/est084.

Goubert C et al. 2015. De novo assembly and annotation of the Asian tiger mosquito (Aedesalbopictus) repeatome with dnaPipeTE from raw genomic reads and comparative analysis with the yellow fever mosquito (Aedes aegypti). Genome Biol. Evol. 7:1192–1205. doi: 10.1093/gbe/evv050.

Hallgren P, Sorita Z, Berglund O, Persson A. 2012. Effects of 17α-ethinylestradiol on individual life-history parameters and estimated population growth rates of the freshwater gastropods Radix balthica and Bithynia tentaculata. Ecotoxicology. 21:803–810. doi: 10.1007/s10646-011-0841-8.

Holt C, Yandell M. 2011. MAKER2: an annotation pipeline and genome- database management tool for second- generation genome projects. BMC Bioinformatics. 12:491. doi: 10.1186/1471-2105-12-491.

Huňová K et al. 2012. Radix spp.: Identification of trematode intermediate hosts in the Czech Republic. Acta Parasitol. 57:273–84. doi: 10.2478/s11686-012-0040-7.

Jarne P, Charlesworth D. 1993. The Evolution of the Selfing Rate in Functionally Hermaphrodite Plants and Animals. Annu. Rev. Ecol. Syst. 24:441–466. http://www.jstor.org/stable/2097186.

Jarne P, Delay B. 1990. Inbreeding depression and self-fertilization in Lymnaea peregra (Gastropoda: Pulmonata). Heredity (Edinb). 64:169–176. doi: 10.1038/hdy.1990.21.

Johansson MP, Ermold F, Kristjánsson BK, Laurila A. 2016. Divergence of gastropod life history in contrasting thermal environments in a geothermal lake. J. Evol. Biol. 1–11. doi: 10.1111/jeb.12928.

Kent WJ. 2002. BLAT — The BLAST -Like Alignment Tool. Genome Res. 12:656–664. doi: 10.1101/gr.229202.

Korf I. 2004. Gene finding in novel genomes. BMC Bioinformatics. 5:59. doi: 10.1186/1471-2105-5-59.

Li H. 2013. Aligning sequence reads, clone sequences and assembly contigs with BWA-MEM. arXiv Prepr. doi: arXiv:1303.3997 [q-bio.GN].

Lomsadze A, Ter-Hovhannisyan V, Chernoff YO, Borodovsky M. 2005. Gene identification in novel eukaryotic genomes by self-training algorithm. Nucleic Acids Res. 33:6494–6506. doi: 10.1093/nar/gki937.

Otto F. 1990. DAPI Staining of Fixed Cells for High-Resolution Flow Cytometly of Nuclear DNA. Methods Cell Biol. 33:105–110. doi: 10.1016/S0091-679X(08)60516-6.

Parra G, Bradnam K, Korf I. 2007. CEGMA: A pipeline to accurately annotate core genes in eukaryotic genomes. Bioinformatics. 23:1061–1067. doi: 10.1093/bioinformatics/btm071.

Patel S, Schell T, Eifert C, Feldmeyer B, Pfenninger M. 2015. Characterizing a hybrid zone between a cryptic species pair of freshwater snails. Mol. Ecol. 24:643–655. doi: 10.1111/mec.13049.

Pfenninger M, Cordellier M, Streit B. 2006. Comparing the efficacy of morphologic and DNA-based taxonomy in the freshwater gastropod genus Radix (Basommatophora, Pulmonata). BMC Evol. Biol. 6:100. doi: 10.1186/1471-2148-6-100.

Pfenninger M, Salinger M, Haun T, Feldmeyer B. 2011. Factors and processes shaping the population structure and distribution of genetic variation across the species range of the freshwater snail radix balthica (Pulmonata, Basommatophora). BMC Evol. Biol. 11:135. doi: 10.1186/1471-2148-11-135.

Quevillon E et al. 2005. InterProScan: Protein domains identifier. Nucleic Acids Res. 33:116–120. doi: 10.1093/nar/gki442.

Quintela M, Johansson MP, Kristjánsson BK, Barreiro R, Laurila A. 2014. AFLPs and Mitochondrial haplotypes reveal local adaptation to extreme thermal environments in a freshwater gastropod. PLoS One. 9. doi: 10.1371/journal.pone.0101821.

R Development Core Team. 2008. R: A language and environment for statistical computing.

Romero PE, Pfenninger M, Kano Y, Klussmann-Kolb A. 2015. Molecular phylogeny of the Ellobiidae (Gastropoda: Panpulmonata) supports independent terrestrial invasions. Mol. Phylogenet. Evol. doi: 10.1016/j.ympev.2015.12.014.

Romero PE, Weigand AM, Pfenninger M. 2016. Positive selection on panpulmonate mitogenomes provide new clues on adaptations to terrestrial life. BMC Evol. Biol. 16. doi: 10.1186/s12862-016-0735-8.

Rosenbaum DM, Rasmussen SGF, Kobilka BK. 2009. The structure and function of G-protein-coupled receptors. Nature. 459:356–363. doi: 10.1038/nature08144.

Rundle SD, Smirthwaite JJ, Colbert MW, Spicer JI. 2011. Predator cues alter the timing of developmental events in gastropod embryos. Biol. Lett. 7:285–287. doi: 10.1098/rsbl.2010.0658.

Simão FA, Waterhouse RM, Ioannidis P, Kriventseva E V., Zdobnov EM. 2015. BUSCO: Assessing genome assembly and annotation completeness with single-copy orthologs. Bioinformatics. 31:3210–3212. doi: 10.1093/bioinformatics/btv351.

Simit A, Hubley R. 2015. RepeatModeler Open-1.0.

Sojo V, Dessimoz C, Pomiankowski A, Lane N. 2016. Membrane proteins are dramatically less conserved than water-soluble proteins across the tree of life. Mol. Biol. Evol. 33:msw164. doi: 10.1093/molbev/msw164.

Sommer U, Adrian R, Bauer B, Winder M. 2012. The response of temperate aquatic ecosystems to global warming: Novel insights from a multidisciplinary project. Mar. Biol. 159:2367–2377. doi: 10.1007/s00227-012-2085-4.

Stanke M, Steinkamp R, Waack S, Morgenstern B. 2004. AUGUSTUS: A web server for gene finding in eukaryotes. Nucleic Acids Res. 32:309–312. doi: 10.1093/nar/gkh379.

Tills O et al. 2011. A genetic basis for intraspecific differences in developmental timing? Evol. Dev. 13:542–548. doi: 10.1111/j.1525-142X.2011.00510.x.

Tills O, Truebano M, Rundle S. 2015. An embryonic transcriptome of the pulmonate snail Radix balthica. Mar. Genomics. 24:259–260. doi: 10.1016/j.margen.2015.07.014.

Wiehn JR, Kopp K, Rezzonico S, Karttunen S, Jokela J. 2002. Family-Level Covariation Between Parasite Resistance and Mating System in a Hermaphroditic Freshwater Snail. Evolution (N. Y). 56:1454–1461. doi: 10.1111/j.0014-3820.2002.tb01457.x.

Xue W et al. 2013. L_RNA_scaffolder: scaffolding genomes with transcripts. BMC Genomics. 14. doi: 10.1186/1471-2164-14-604.

Yu TL, Deng YH, Zhang J, Duan LP. 2016. Size-assortative copulation in the simultaneously hermaphroditic pond snail Radix auricularia (Gastropoda: Pulmonata). Anim. Biol. doi: 10.1163/15707563-00002501.

Zdobnov EM, Apweiler R. 2001. InterProScan - an integration platform for the signature-recognition methods in InterPro. Bioinformatics. 17:847–848. doi: 10.1093/bioinformatics/17.9.847.

